# Development of a High-Throughput TR-FRET Assay for Identification of Small Molecule Inhibitors of the LILRB4 (ILT3)-SCG2 Immune Checkpoint Interaction

**DOI:** 10.64898/2026.05.19.726368

**Authors:** Somaya A. Abdel-Rahman, Moustafa T. Gabr

**Affiliations:** Department of Radiology, Molecular Imaging Innovations Institute (MI3), Weill Cornell Medicine, New York, NY 10065, USA

**Keywords:** TR-FRET, Small molecules, High-throughput screening, LILRB4, ILT3

## Abstract

Leukocyte immunoglobulin-like receptor B4 (LILRB4, ILT3) is an inhibitory immune checkpoint expressed on myeloid cells, where it contributes to immunosuppression within the tumor microenvironment. Secretogranin 2 (SCG2) has recently been identified as a functional ligand of LILRB4, yet small molecule modulators of this interaction remain unexplored. Here, we report the development of a high-throughput time-resolved fluorescence resonance energy transfer (TR-FRET) assay to interrogate the LILRB4 (ILT3)-SCG2 interaction. The assay demonstrated robust performance and was validated using a blocking anti-LILRB4 antibody, consistent with orthogonal ELISA measurements. Pilot screening of chemical libraries identified 23 primary hits, of which two compounds, BMS-813160 and PSB-603, showed reproducible, dose-dependent inhibition with TR-FRET IC_50_ values of 26.7 ± 1.03 µM and 37.2 ± 2.14 µM, respectively. Activity was confirmed by ELISA, supporting the robustness of the assay. This platform enables high-throughput discovery of first-in-class small molecule modulators of the LILRB4-SCG2 immune checkpoint and provides a foundation for targeting myeloid-driven immunosuppression.

## Introduction

Immune checkpoint blockade (ICB) has transformed the therapeutic landscape of cancer by restoring antitumor immune responses, leading to durable clinical benefit in a subset of patients.^1-3^ Monoclonal antibodies targeting T-cell inhibitory receptors such as programmed cell death protein 1 (PD-1), its ligand PD-L1, and cytotoxic T-lymphocyte-associated protein 4 (CTLA-4) have demonstrated remarkable efficacy across multiple malignancies.^3,4^ However, despite these advances, the majority of patients either fail to respond or develop resistance due to complex immunosuppressive mechanisms within the tumor microenvironment (TME).^5-7^ These limitations have driven increasing interest in identifying alternative immune checkpoints and complementary pathways that contribute to immune evasion.

While most clinically approved immune checkpoint therapies focus on T-cell-intrinsic mechanisms, there is growing recognition that myeloid cells play a central role in shaping immunosuppressive TMEs.^8-10^ Myeloid-derived suppressor cells (MDSCs), tumor-associated macrophages (TAMs), and tolerogenic dendritic cells contribute to suppression of cytotoxic T-cell responses, promotion of tumor growth, and resistance to immunotherapy.^9,11^ Consequently, targeting myeloid immune checkpoints has emerged as a promising strategy to overcome resistance and enhance the efficacy of existing immunotherapies.

Leukocyte immunoglobulin-like receptor B4 (LILRB4), also known as ILT3, is a type I transmembrane inhibitory receptor predominantly expressed on monocytic cells, including MDSCs and dendritic cells.^12-14^ LILRB4 contains intracellular immunoreceptor tyrosine-based inhibitory motifs (ITIMs) that recruit phosphatases such as SHP1 and SHP2 upon activation, leading to suppression of immune signaling pathways.^15-17^

Elevated expression of LILRB4 has been associated with immunosuppressive phenotypes, tumor progression, and poor clinical outcomes in several cancers.^16-18^ Importantly, recent studies have demonstrated that LILRB4 signaling contributes to suppression of T-cell activation and promotes immune evasion, highlighting its potential as a therapeutic target in cancer immunotherapy.

A major advance in understanding LILRB4 biology came with the identification of secretogranin 2 (SCG2) as a functional ligand of LILRB4.^19^ SCG2, a member of the granin family of secreted proteins, is expressed in endocrine and neuronal tissues and has been implicated in diverse biological processes, including neuroinflammation and immune regulation.^20,21^ Recent work has demonstrated that secretogranin 2 (SCG2) binds specifically to the extracellular domain of LILRB4, triggering receptor phosphorylation, recruitment of SHP1/SHP2, and activation of downstream signaling pathways, including STAT3.^19^ This interaction promotes monocytic immunosuppression, enhances MDSC function, and supports tumor growth in a T-cell-dependent manner.^19^ These findings establish the LILRB4-SCG2 axis as a novel hormone-checkpoint pathway that orchestrates immunosuppressive signaling within the TME.

Despite the therapeutic promise of targeting immune checkpoints, current approaches rely predominantly on monoclonal antibodies, which present several limitations, including high production costs, limited tissue penetration, and restricted pharmacokinetic flexibility.^22-24^ In contrast, small molecules offer several advantages, including oral bioavailability, improved tumor penetration, and the ability to fine-tune exposure through dosing strategies.^23,25^ However, the development of small molecule inhibitors targeting protein-protein interactions (PPIs), such as LILRB4-SCG2, remains challenging due to the typically large and shallow binding interfaces involved. To date, no small molecule inhibitors of the LILRB4-SCG2 interaction have been reported, highlighting a significant unmet need in the field.

High-throughput screening (HTS) technologies are essential for the discovery of small molecule modulators of PPIs. Among these, time-resolved fluorescence resonance energy transfer (TR-FRET) has emerged as a powerful and widely used platform due to its homogeneous format, high sensitivity, low background, and compatibility with miniaturized screening formats. As we have shown in our previous work,^26-31^ TR-FRET assays enable quantitative measurement of PPIs by detecting proximity-dependent energy transfer between donor and acceptor fluorophores, providing a robust readout for screening chemical libraries.

In this study, we report the development and optimization of a TR-FRET-based high-throughput assay for the LILRB4 (ILT3)-SCG2 interaction. Building on our previous work in immune checkpoint assay development,^26-31^ we established a robust and reproducible platform suitable for screening small molecule libraries. The assay was validated using a LILRB4-blocking antibody and benchmarked against orthogonal assays. We further demonstrate the utility of this platform through pilot screening of chemical libraries and identification of small molecule hits. This work provides a critical enabling tool for discovery of first-in-class small molecule modulators of the LILRB4-SCG2 immune checkpoint and supports future efforts targeting myeloid-driven immunosuppression in cancer.

## Methods

The overall workflow for development of the TR-FRET assay designed to monitor the interaction between LILRB4 (ILT3) and SCG2 is depicted in Figure 1. Recombinant human SCG2 carrying C-terminal polyhistidine tag was purchased from SinoBiological (catalog number 13441-H08H), while the extracellular domain (ECD) of human LILRB4 (ILT3) fused to the Fc region of human IgG1 was obtained from SinoBiological (catalog number 16742-H02H). Detection of the interacting proteins was achieved using a terbium cryptate-labeled anti-His monoclonal antibody as the donor fluorophore and an XL665-conjugated anti-human Fc antibody as the acceptor, both sourced from Revvity (catalog numbers 61HISTLF and 61HFCXLF, respectively). Assay optimization and validation studies, including protein titration, donor-to-acceptor ratio optimization, and incubation time evaluation, are presented in Figures S1-S4 and Tables S1-S2.

**Figure 1.**
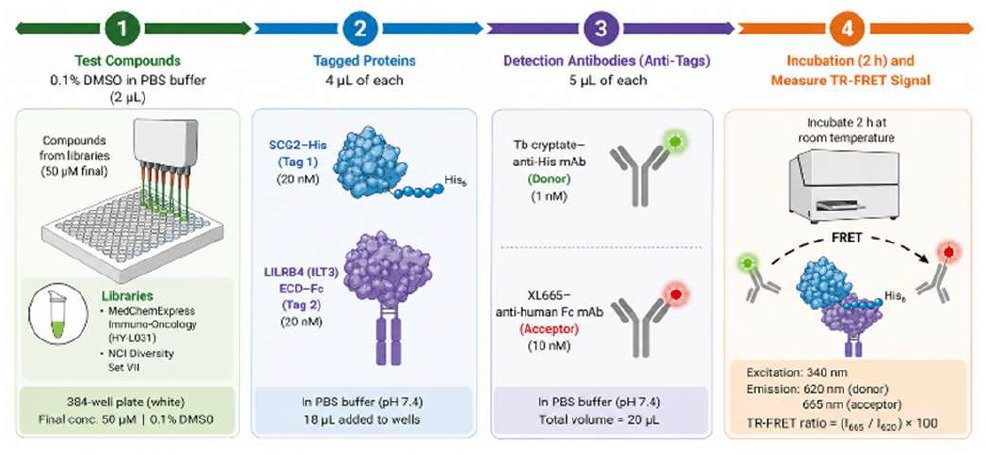
Our established workflow for LILRB4-SCG2 TR-FRET assay.

TR-FRET measurements were performed using a Tecan Spark plate reader. The donor emission was recorded at 620 nm following excitation at 340 nm, and acceptor emission was measured at 665 nm under identical excitation conditions. Data acquisition parameters included 50 flashes per well, a 50 µs delay time, and a 400 µs integration window. The assay reaction mixture was freshly prepared immediately prior to use and consisted of LILRB4-ECD-Fc at a final concentration of [20 nM], SCG2-His at [20 nM], terbium-labeled anti-His antibody at [1 nM], and XL665-labeled anti-Fc antibody at [10 nM] in phosphate-buffered saline (PBS, pH 7.4).

Compounds from both chemical libraries (Small molecule Immuno-Oncology Compound Library from MedChemExpress (catalog number HY-L031) and the National Cancer Institute (NCI) Diversity Set VII chemical library) were prepared as stock solutions in DMSO and transferred to medium-binding white 384-well plates (Greiner, #784075), resulting in a final compound concentration of [50 µM] with a DMSO content of 0.1% (v/v). Following compound dispensing (2 µL per well), the assay mixture (18 µL) was added, and plates were incubated for 2 hours at room temperature to allow equilibrium binding. After incubation, fluorescence signals were measured as described above. The TR-FRET response was expressed as the ratio of acceptor to donor emission, calculated as (I_665_ I_620_) × 100.

Each assay plate included multiple control wells to ensure data quality and reproducibility. Negative control wells contained DMSO vehicle alone, while background control wells lacked one of the tagged assay components to account for nonspecific signal. A blocking anti-LILRB4 antibody (h128-3, catalog number HV126013) was included to define maximal inhibition. Percent inhibition values were normalized using plate-specific controls, with 0% corresponding to the DMSO condition and 100% corresponding to the positive control. Assay performance was assessed using the Z’ factor, calculated based on the interquartile mean and variability of control wells.^32^

For hit selection, compounds producing ≥50% reduction in the TR-FRET signal relative to the negative control were classified as primary hits. To exclude false positives arising from optical interference, donor channel fluorescence (620 nm) was analyzed independently. Compounds that produced significant changes in donor emission consistent with signal quenching or fluorescence artifacts were considered assay interferents and excluded from further analysis. Dose-response experiments were performed with n = 5 replicates, and data are reported as mean ± standard deviation.

An enzyme-linked immunosorbent assay (ELISA) was used as an orthogonal method to validate inhibition of the LILRB4 (ILT3)- SCG2 interaction. High-binding 96-well plates were coated with recombinant human LILRB4 (ILT3) ECD-Fc (2 µg/mL) in phosphate-buffered saline (PBS) and incubated overnight at 4 °C. Plates were washed with PBS containing 0.05% Tween-20 (PBST) and blocked with 3% bovine serum albumin (BSA) in PBS for 1 h at room temperature.

Test compounds were prepared in assay buffer (PBS containing 0.1% BSA and 0.1% DMSO) and pre-incubated with His-tagged SCG2 (100 nM) for 30 min at room temperature. The mixture was then added to LILRB4-coated plates and incubated for 1 h to allow binding. Following incubation, plates were washed with PBST and bound SCG2 was detected using an anti-His horseradish peroxidase (HRP)-conjugated antibody. Signal was developed using tetramethylbenzidine (TMB) substrate and quenched with 1 M H2SO4. Absorbance was measured at 450 nm using a plate reader. Data were normalized and expressed as percent inhibition relative to DMSO-treated controls (0% inhibition) and maximal inhibition defined by a blocking anti-LILRB4 antibody (100% inhibition). Dose-response curves were generated and IC_50_ values were calculated using nonlinear regression analysis (GraphPad Prism v10).

## Results and discussion

As part of our ongoing efforts to develop screening platforms for discovery of small molecule modulators targeting PPIs,^26-31^ we established a TR-FRET-based assay to interrogate the interaction between LILRB4 (ILT3) and its ligand SCG2 (Figure 2). TR-FRET relies on a pair of fluorophores (a donor and an acceptor) that undergo non-radiative energy transfer when brought into close spatial proximity. In the context of a PPI assay, each binding partner is either directly or indirectly labeled with one of these fluorophores. Upon interaction of the two proteins, excitation of the donor results in energy transfer to the acceptor, producing a signal proportional to the extent of binding.^31^ Disruption of the interaction by inhibitory compounds leads to a reduction in the TR-FRET signal, providing a quantitative readout suitable for screening applications.^31^

**Figure 2.**
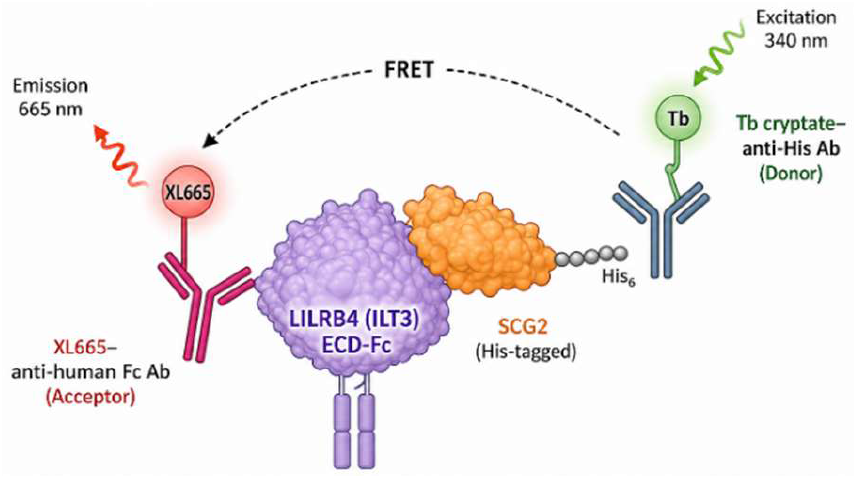
Schematic representation of the TR-FRET assay developed for identification of small molecule inhibitors of the LILRB4 (ILT3)-SCG2 interaction.

To establish optimal assay conditions for the LILRB4-SCG2 interaction, we systematically evaluated multiple combinations of tagging strategies and donor-acceptor detection pairs. Cross-titration experiments were performed to assess signal generation across a range of protein concentrations and labeling configurations. In parallel, assay conditions were adapted to a 384-well plate format to enable high-throughput compatibility. Additional parameters, including buffer composition, plate type, reagent addition sequence, and incubation time, were optimized to maximize assay performance (see Figures S1-S4 and Tables S1-S2 for optimization efforts). The most robust signal was obtained using terbium cryptate-labeled anti-His antibody as the donor and XL665-labeled anti-human Fc antibody as the acceptor, with an optimal donor-to-acceptor ratio of 1:10. Under these conditions, the assay exhibited a signal-to-background (S/B) ratio of 9.67, with optimal protein concentrations of 20 nM for both SCG2-His and LILRB4-ECD-Fc.

To further characterize assay performance, we evaluated the dependence of the TR-FRET signal on acceptor concentration. Increasing levels of XL665-labeled detection antibody resulted in a hyperbolic increase in the TR-FRET ratio, consistent with efficient energy transfer upon complex formation (Figure 3A). The ability of the assay to detect disruption of the LILRB4-SCG2 interaction was assessed using a blocking anti-LILRB4 antibody (h128-3). A clear dose-dependent decrease in TR-FRET signal was observed with increasing concentrations of the antibody (Figure 3B), confirming that the assay accurately reports on target engagement. Analysis of the dose-response curve yielded an IC_50_ value of 6.71 ± 0.45 nM. To validate these findings, the inhibitory activity of the antibody was independently evaluated using in-house ELISA-based assay, which produced a comparable IC_50_ value of 9.56 ± 0.38 nM.

**Figure 3.**
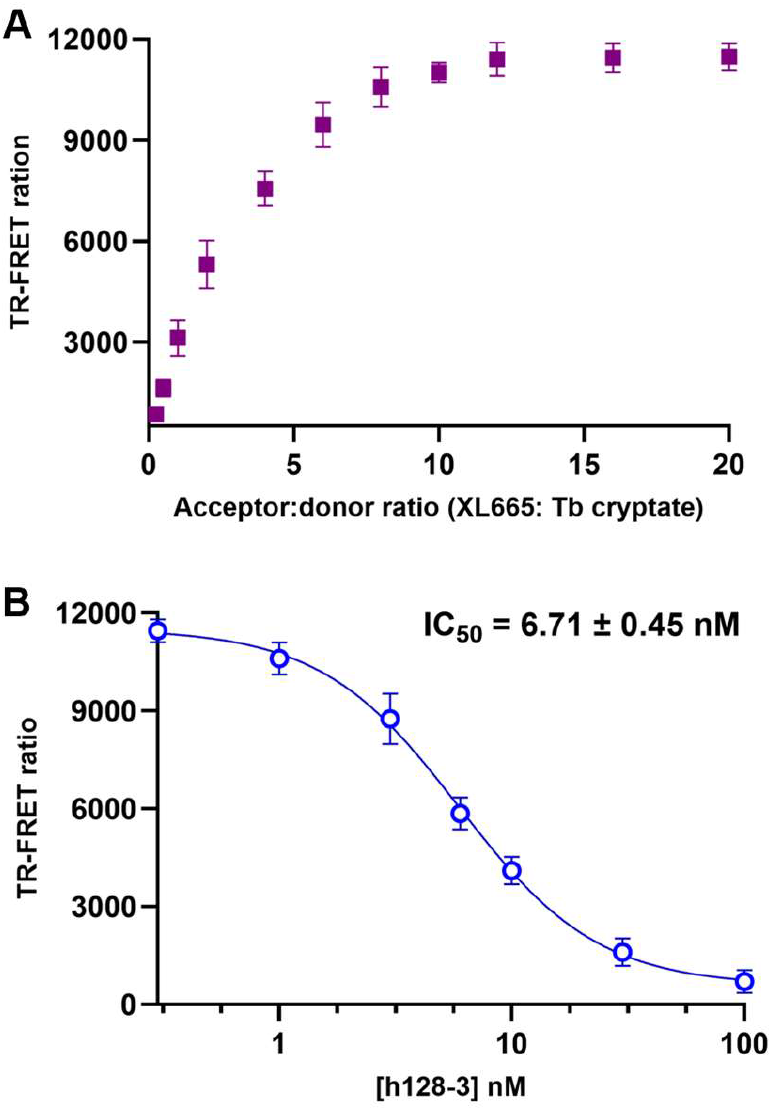
**(A)** TR-FRET signal of the LILRB4 (ILT3)-SCG2 interaction assay as a function of acceptor concentration. **(B)** Dose-response curve of anti-LILRB4 antibody in the TR-FRET assay. Error bars represent standard deviation (n = 5).

The strong agreement between TR-FRET and ELISA-based measurements supports the reliability of the assay for quantifying disruption of the LILRB4-SCG2 interaction. Assessment of assay robustness demonstrated a Z’ factor of 0.73 ± 0.08, indicating excellent suitability for HTS applications. The Z’ factor reflects the separation between positive and negative controls relative to signal variability and is a widely accepted metric for assay quality.^32^ In addition, evaluation of DMSO tolerance revealed that assay performance remained stable up to 3.0% (v/v), confirming compatibility with standard compound screening conditions.

Following optimization and validation, we applied the assay to a pilot screening campaign using a subset of the National Cancer Institute (NCI) Diversity Set VII as well as the Immuno-Oncology Compound Library from MedChemExpress (catalog number HY-L031) (Figures 4A and 4B). Compounds were screened at a final concentration of 50 µM, and inhibition of the TR-FRET signal was used as the primary readout. This initial screening identified 8 compounds from the NCI library and 15 compounds from the MedChemExpress library that exhibited ≥50% inhibition. As the primary objective of this screen was to demonstrate assay feasibility, these hits were evaluated at a preliminary level and not pursued further due to limited suitability for downstream medicinal chemistry optimization.

**Figure 4.**
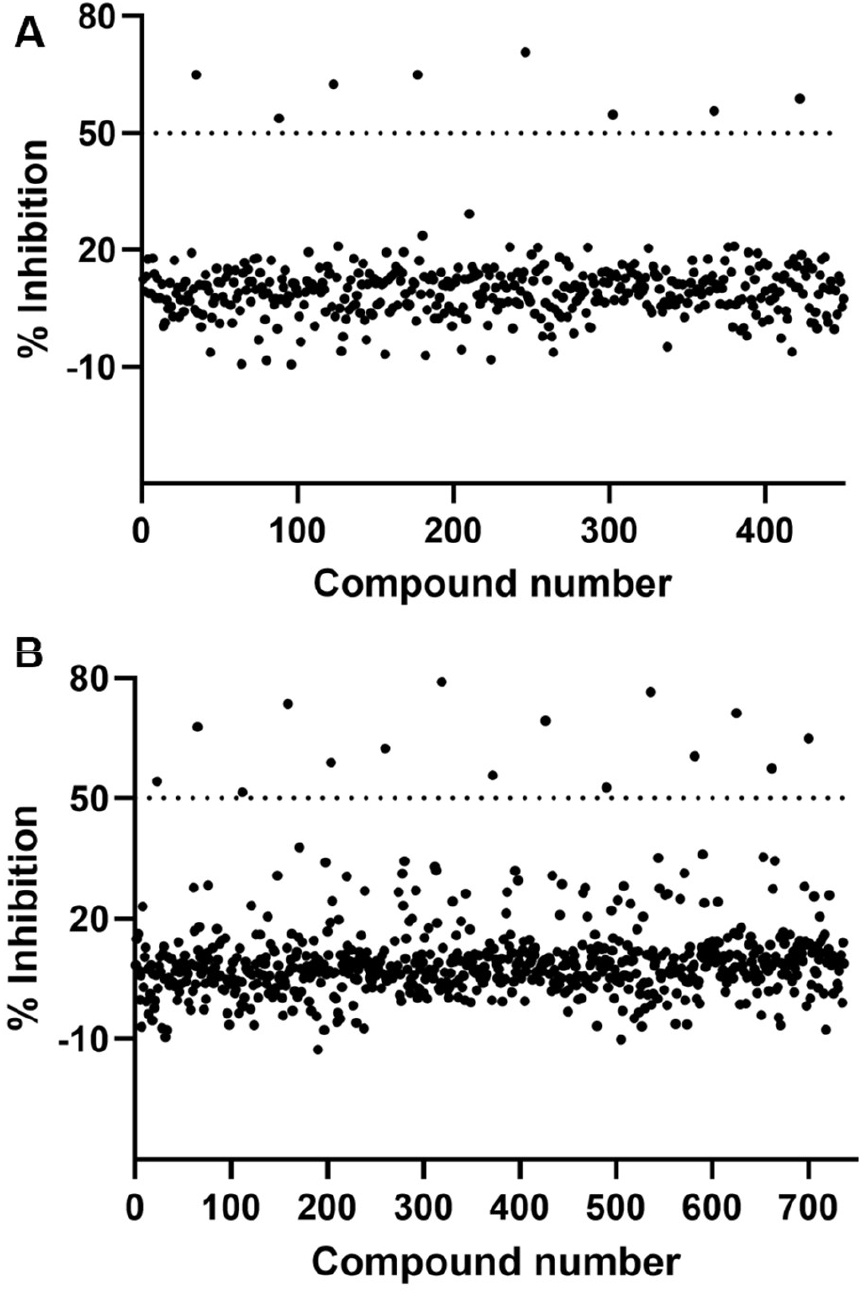
**(A)** Scatter plot of single-dose screening data from the NCI Diversity Set VII. **(B)** Scatter plot of screening data from the MedChemExpress immuno-oncology library.

Following identification of the primary hits from the pilot screening campaign, we performed secondary evaluation to assess reproducibility and dose-dependent inhibition. A total of 23 compounds meeting the ≥50% inhibition threshold, including 8 compounds from the NCI Diversity Set VII and 15 compounds from the MedChemExpress Immuno-Oncology library, were selected for follow-up analysis. These compounds were retested in dose-response format using the LILRB4 (ILT3)-SCG2 TR-FRET assay.

Among the tested compounds, BMS-813160 and PSB-603 consistently demonstrated reproducible, dose-dependent inhibition of the LILRB4 (ILT3)-SCG2 interaction. The chemical structures of these validated hits are shown in Figure 5A. BMS-813160 inhibited the interaction with a TR-FRET IC_50_ value of 26.7 ± 1.03 µM, while PSB-603 showed an IC_50_ value of 37.2 ± 2.14 µM (Figures 5B,C). The remaining primary hits either failed to reproduce the initial screening activity or did not show clear dose-dependent inhibition. To further validate these findings, BMS-813160 and PSB-603 were evaluated using an orthogonal ELISA-based LILRB4-SCG2 inhibition assay. Consistent with the TR-FRET results, BMS-813160 and PSB-603 inhibited the interaction with ELISA IC_50_ values of 20.2 ± 1.59 µM and 42.7 ± 3.44 µM, respectively (Figures 5D,E). The agreement between TR-FRET and ELISA results supports the reproducibility of these compounds as validated small molecule modulators of the LILRB4 (ILT3)-SCG2 interaction.

**Figure 5.**
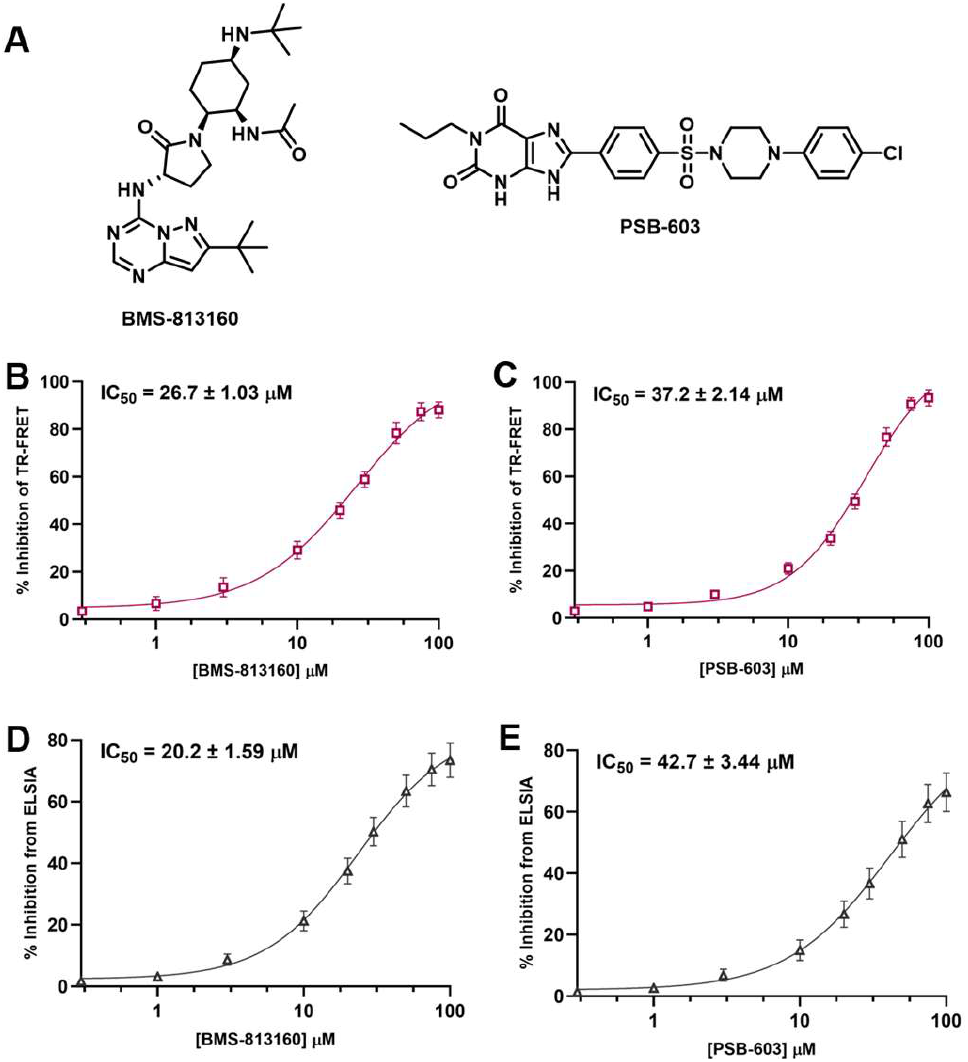
**(A)** Chemical structures of the validated hits BMS-813160 and PSB-603. **(B-C)** Dose–response curves obtained from the TR-FRET assay showing inhibition of the LILRB4 (ILT3)- SCG2 interaction by BMS-813160 and PSB-603. **(D-E)** Orthogonal validation using ELISA demonstrating comparable inhibitory activity. Data represent mean ± SD (n = 5).

In summary, we developed and validated a robust TR-FRET-based high-throughput assay for quantitative interrogation of the LILRB4 (ILT3)-SCG2 PPI. The assay demonstrates strong performance characteristics, including a high signal-to-background ratio, excellent Z’ factor, and compatibility with standard HTS conditions. Importantly, the assay accurately reports disruption of the LILRB4-SCG2 interaction, as demonstrated by dose-dependent inhibition with a blocking antibody and strong agreement with orthogonal ELISA measurements.

Application of the assay to pilot screening campaigns enabled identification of small molecule modulators of this interaction. Secondary validation revealed two reproducible inhibitors, BMS-813160 and PSB-603, which exhibited dose-dependent activity in both TR-FRET and ELISA assays. Although these compounds display moderate potency, their consistent activity across orthogonal platforms highlights the tractability of the LILRB4-SCG2 interaction for small molecule targeting. Collectively, this work establishes a scalable and reliable screening platform for discovery of small molecule inhibitors of the LILRB4-SCG2 immune checkpoint. These findings provide a foundation for future medicinal chemistry efforts aimed at developing first-in-class therapeutics targeting myeloid-driven immunosuppression and may support combination strategies with existing immune checkpoint therapies.

