## Supporting Information for "Development of a High-Throughput TR-FRET Assay for Identification of Small Molecule Inhibitors of the LILRB4 (ILT3)-SCG2 Immune Checkpoint Interaction"

*Electronic Supplementary Information*

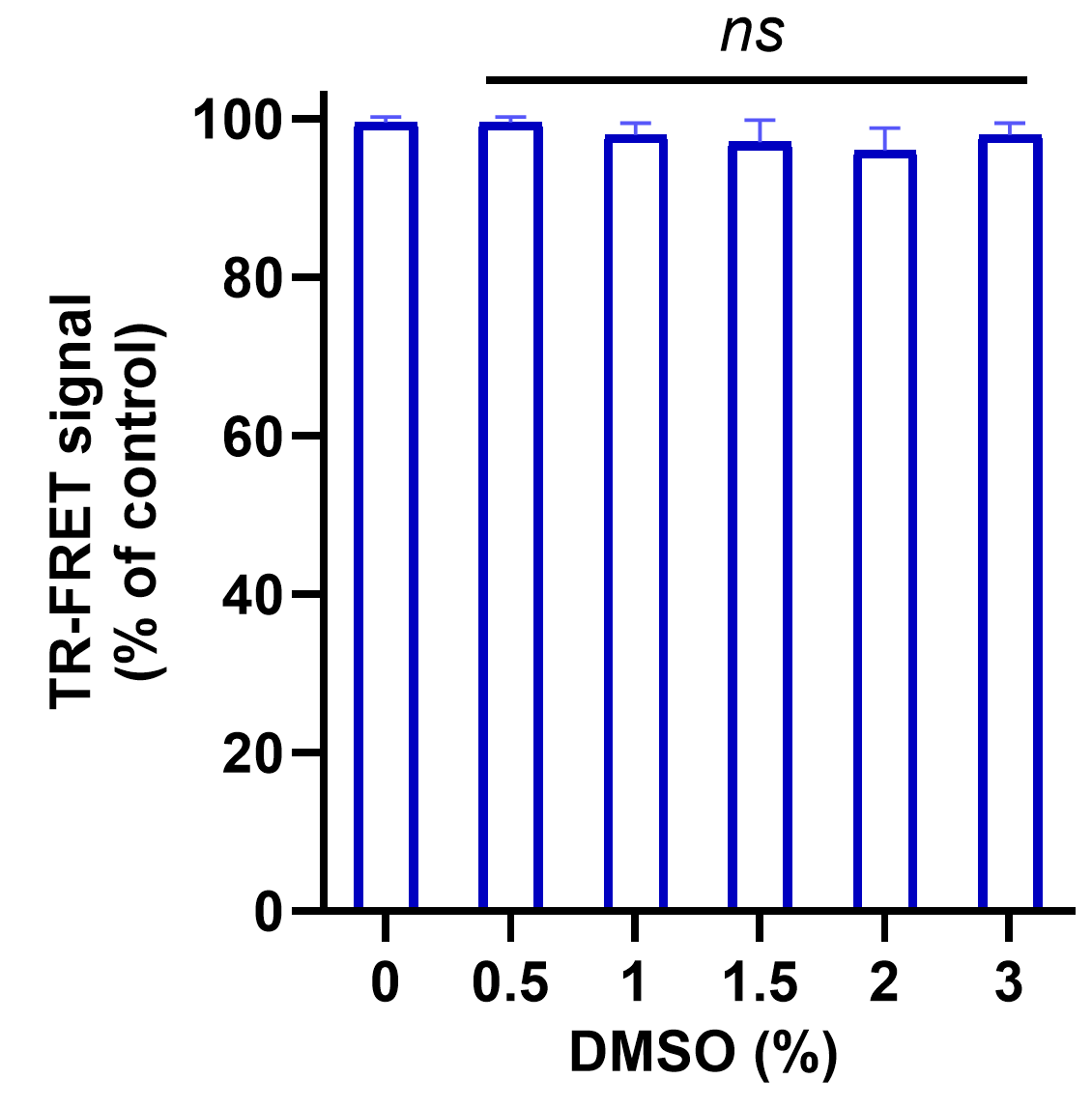

**Fig. S1.** DMSO tolerance of the developed LILRB4/SCG2 TR-FRET assay in the presence of various DMSO (%). Error bars represent standard deviation (n = 5). *ns* denotes non-significant .

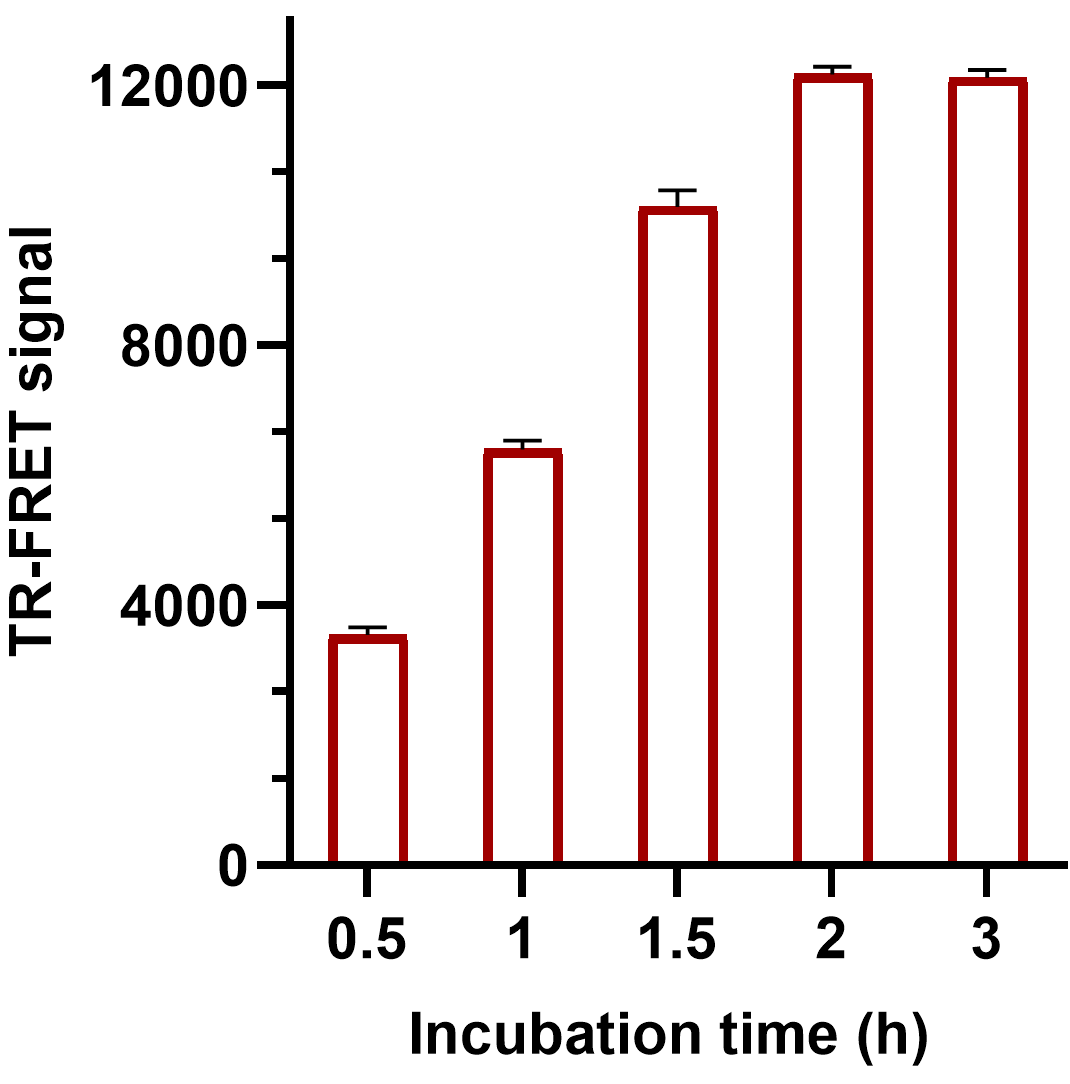

**Fig. S2.** The impact of various incubation times (h) on TR-FRET ratio in the LILRB4/SCG2 TR-FRET assay. Error bars represent standard deviation (n = 5).

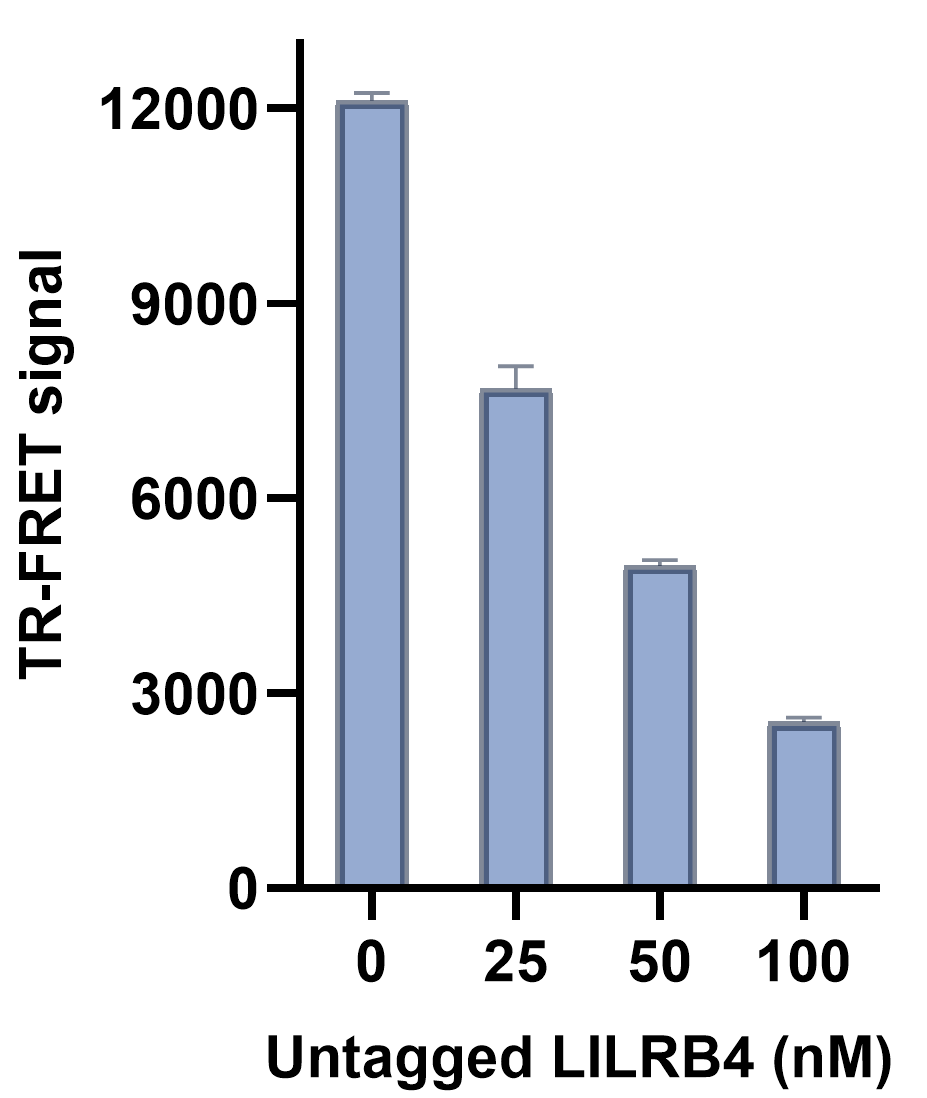

**Fig. S3.** Dose-dependent reduction in the TR-FRET signal of the LILRB4/SCG2 upon incubation with increasing concentrations of untagged LILRB4. Error bars represent standard deviation (n = 5).

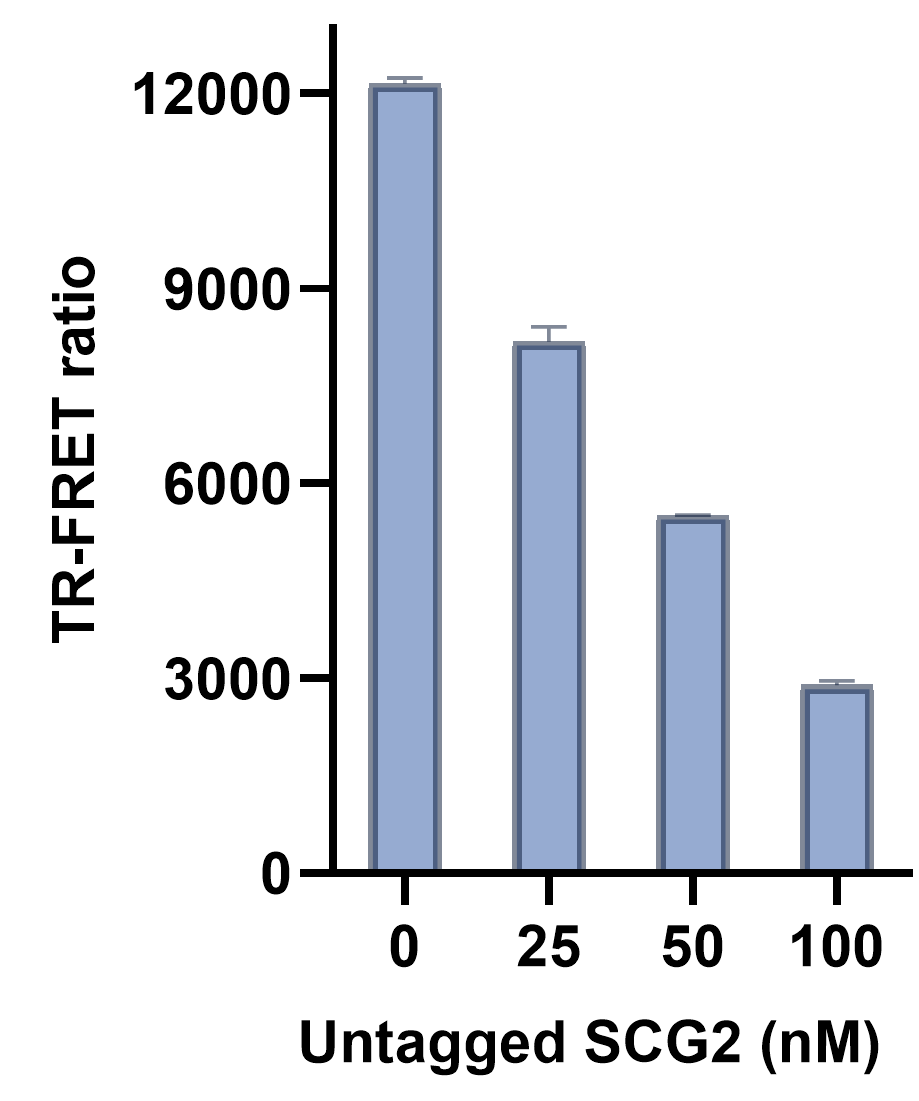

**Fig. S4.** Dose-dependent reduction in the TR-FRET signal of the LILRB4/SCG2 upon incubation with increasing concentrations of untagged SCG2. Error bars represent standard deviation (n = 5).

**Table S1.** 2D titration of LILRB4 and SCG2 in the presence of Tb cryptate labeled anti-His mAb (1 nM), and XL665 labeled anti-human Ab (10 nM) and the corresponding signal-to-background ratio from the TR-FRET assay.

|  | **300 nM LILRB4** | **200 nM LILRB4** | **100 nM LILRB4** | **50 nM LILRB4** | **20 nM LILRB4** | **10 nM LILRB4** | **1 nM LILRB4** |
| --- | --- | --- | --- | --- | --- | --- | --- |
| **300 nM SCG2** | 9.8 | 9.9 | 9.7 | 9.4 | 9.5 | 5.1 | 2.7 |
| **200 nM SCG2** | 9.9 | 10.0 | 9.8 | 9.5 | 9.6 | 4.8 | 2.5 |
| **100 nM SCG2** | 9.7 | 9.8 | 9.6 | 9.3 | 9.5 | 4.5 | 2.3 |
| **50 nM SCG2** | 9.4 | 9.5 | 9.4 | 9.1 | 9.6 | 4.0 | 2.0 |
| **20 nM SCG2** | 9.3 | 9.4 | 9.5 | 9.6 | **9.67** | 3.9 | 1.9 |
| **10 nM SCG2** | 6.2 | 6.5 | 6.8 | 6.9 | 6.7 | 3.8 | 1.6 |
| **1 nM SCG2** | 2.2 | 2.3 | 2.4 | 2.5 | 2.3 | 1.8 | 1.2 |

**Table S2.** The impact of different buffer compositions on the signal-to-background ratio from the LILRB4/SCG2 TR-FRET assay.

| **Buffer** | **Signal-to-background ratio** |
| --- | --- |
| Tris buffer, pH 7.0 | 6.35 |
| Tris buffer, pH 7.0 + DTT | 6.68 |
| Tris buffer, pH 8.0 | 6.12 |
| HEPES buffer, pH 7.5 | 8.14 |
| HEPES buffer, pH 7.5 + DTT | 7.25 |
| HEPES buffer, pH 8.0 | 6.24 |
| PBS, pH 7.4 | 9.67 |
| PBS, pH 7.4 + DTT | 9.13 |
